# Genes shielded, repeats exposed: mutation bias in the midge *Chironomus riparius*

**DOI:** 10.64898/2025.12.01.691625

**Authors:** María Esther Nieto-Blázquez, Markus Pfenninger

## Abstract

Mutation is the fundamental source of genetic variation, yet growing evidence shows that mutations are not uniformly distributed across genomes but are shaped by genomic architecture, DNA-repair dynamics, and environmental conditions. Here, we investigate fine-scale determinants of mutation distribution in the non-biting midge *Chironomus riparius*, an ecologically important freshwater insect widely used in ecotoxicology. We integrated mutation data from five independent studies, including spontaneous mutation-accumulation experiments and multigenerational exposure assays involving cadmium, benzo[a]pyrene, tyre and road wear particles, and varying generational time. In total, we analysed 420 single-nucleotide mutations mapped to the chromosome-scale *C. riparius* reference genome.

Using a Bayesian modelling framework, we tested whether mutation density is (i) randomly distributed, (ii) non-randomly distributed, or predicted by (iii) distance to telomeres and centromeres, (iv) proximity to genes, or (v) distance to repetitive elements. Models were compared using leave-one-out cross-validation (LOO-CV). We also quantified the proportion of mutations in exons and evaluated the synonymous vs. non-synonymous spectrum using BayesFactor in R.

The best-supported model incorporated non-linear effects of genomic position and distance to genes, identifying proximity to coding regions as the dominant predictor of mutation rate. Mutation density increased with distance from genes, indicating strong protection of genic regions. A model including repetitive elements showed nearly equivalent support, suggesting that functional and structural features jointly shape mutational landscapes. Only 9.8% of mutations occurred in exons despite exons representing 22.85% of callable sites, demonstrating marked depletion of exonic mutations. Among exonic mutations, 70.7% were non-synonymous—statistically indistinguishable from the neutral expectation (75%).

These findings show that mutation processes in *C. riparius* are strongly structured by genome architecture, with implications for evolutionary genomics, ecotoxicology, and population-genetic inference.

## Introduction

Mutation, the process that generates permanent alterations in the sequence of nucleic acids, is the essential raw material for evolution by natural selection (Lynch 2010). Historically, mutation was often treated in population genetics models as a random process, occurring uniformly across the genome and being independent of genomic location or underlying sequence structure. This “random walk” model provided a simple framework for estimating overall mutation rates and interpreting genetic drift (Scally and Durbin 2012).

While population genomic data reveal only mutations that persist after selection, mutation-accumulation and exposure experiments target the primary generation of mutations prior to selective filtering, *de novo* mutations. Traditionally, mutations were assumed to occur randomly and independently along the genome—a central assumption of many evolutionary and population-genetic models. However, high-resolution sequencing studies and advances in comparative genomics have decisively overturned this view. It is now well-established that the rate of mutation is fundamentally non-uniform; mutation rates vary extensively across evolutionary timescales, among taxa, and crucially, within the genome of a single organism (Hodgkinson and Eyre-Walker 2011). This variation, often referred to as mutation rate heterogeneity (MRH), defines a genomic landscape characterized by mutational hotspots— regions with elevated mutation rates—and coldspots—regions where mutations are conspicuously rare (Liu et al. 2017; Nesta et al. 2021). Accumulating evidence across taxa indicates additional deviations from randomness, including clustering of mutations in specific genomic regions, strand asymmetries, and context-dependent mutation probabilities (Harris 2015; Smith et al. 2018). Understanding the mechanisms underlying these patterns—such as DNA-repair efficiency, replication timing, and sequence context—has both theoretical and applied implications. From a theoretical standpoint, it refines models of molecular evolution and genomic architecture; from an applied perspective, identifying regions disproportionately affected by environmental mutagens is critical for predicting the evolutionary consequences of pollution and for integrating mutation data into environmental risk assessment frameworks (Dearfield and Moore 2005; Heflich et al. 2020; Portugal et al. 2022).

The study of MRH has shifted from simply acknowledging non-randomness to actively identifying the structural and functional correlates that impose profound biases on mutation accumulation. These intra-genomic biases are not random noise; they reflect the complex, often conflicting, outcomes of local DNA damage generation, the efficiency of cellular repair machinery, and the selective pressures unique to specific genomic locations (Chuang and Li 2004; Poetsch et al. 2018; Volkova et al. 2020; Monroe et al. 2022; Majic and Payne 2023; Hara and Kuraku 2025). The non-random distribution of mutations across a eukaryotic genome is thus primarily driven by the underlying structural and functional architecture, making the location of a mutation relative to key genomic features a powerful predictor of its likelihood (Seplyarskiy et al. 2012). To accurately map and model the fine-scale mutational landscape, it is critical to assess the quantitative influence of major genomic features.

The relationship between mutation rate and gene location is one of the most studied and mechanistically complex predictors of MRH. In many model systems, transcribed regions— especially those with high expression—exhibit a lower mutation rate than surrounding non-coding DNA. This protective effect is largely attributed to the process of Transcription-Coupled DNA Repair (TCR), where active transcription machinery effectively flags DNA lesions in the transcribed strand, leading to expedited repair and therefore fewer permanent mutations (Deger et al. 2019; Törmä et al. 2020). The proximity of a mutation to a gene is thus a critical proxy for the local repair environment. Mutations occurring immediately adjacent to or within a coding sequence are often subject to stronger repair pressures and functional constraints. Conversely, mutations occurring far from genes, in vast intergenic regions, may be subject to less efficient general repair mechanisms, leading to a potentially higher local mutation rate. However, this relationship is not universal; in some contexts, the act of frequent transcription can increase the accessibility of the DNA, or the formation of structures like R-loops can lead to increased damage, creating mutation hotspots around highly active genes (Polak et al. 2014).

The architecture of the chromosome itself significantly impacts MRH. Telomeres (chromosome ends) and centromeres (the constricted region critical for cell division) are major structural elements characterized by large tracts of highly compacted, non-coding, heterochromatic DNA (Bloom 2014). These regions are widely recognized as sites of genomic instability. Centromeres and telomeres are often late-replicating and are associated with a less efficient or slower DNA repair environment compared to gene-rich, euchromatic regions (Han et al. 2016). The dense, repetitive nature of these regions makes them prone to double-strand break (DSB) formation, and the errors introduced during the non-homologous end joining (NHEJ) and homologous recombination (HR) processes often lead to localized mutation clusters (Cannan and Pederson 2016; Hussmann et al. 2021; Miller et al. 2023). Consequently, mutations are frequently found to accumulate at a higher rate near the physical termini (telomeres) and the central structural regions (centromeres) of chromosomes.

The eukaryotic genome is replete with repetitive elements, including simple sequence repeats (SSRs), microsatellites, and transposable elements (TEs). Although often labeled “junk DNA,” these elements are highly dynamic and constitute significant mutation drivers (Ellegren 2004). Repetitive sequences are inherently unstable due to the mechanisms of DNA replication and repair. They are prone to replication errors, primarily strand slippage (or replication-slippage), which leads to insertions and deletions (indels) of the repeat units, thereby acting as localized mutation hotspots (Bzymek and Lovett 2001; Cabral-de-Mello and Palacios-Gimenez 2025). Furthermore, the density and location of TEs can indirectly affect mutation rates by facilitating ectopic recombination or by disrupting adjacent chromatin structures. The presence of a high density of repetitive elements is therefore expected to correlate with an elevated local mutation rate due to inherent difficulties in maintaining sequence fidelity in these regions (Bzymek and Lovett 2001).

The non-biting midge, *Chironomus riparius* (Diptera: Chironomidae), served as the model organism for the present study. *C. riparius* is a keystone species in freshwater ecosystems and is a standard indicator species in ecotoxicology due to its sensitivity and wide geographic distribution (Pinder 1986). The aquatic environment subjects this organism to chronic exposure to genotoxic stress mutagenic agents, including heavy metals and pollutants, which are known to induce high levels of DNA damage (Doria et al. 2021; Pinto et al. 2024), and may also affect germline integrity.

We focus on the initial occurrence of *de novo* mutations using experimental approaches that minimize selective filtering. The central rationale of this study is to move beyond the assumption of random mutation by testing the contribution of underlying genomic architecture to the distribution of mutations in *C. riparius* by integrating available mutation-location data. Our study aims to: 1) test for and model the non-random distribution of the local mutation rate across the *C. riparius* genome. We hypothesize that mutation distribution is non-random and can be significantly predicted by the distance to genomic features (i.e genes, proximity to telomeres/centromeres, and density of repetitive elements); and 2) test for functional enrichment within the identified mutation hotspots. We hypothesize that a mutation occurring randomly and neutrally is expected to result in a non-synonymous (amino acid changing) change approximately 75% of the time (Nei and Gojobori 1986). By analyzing how architectural features govern mutation distribution and by confirming the functional significance of these mutations through an enrichment test, this study provides novel and quantitative insights into the forces shaping the genome and thus the evolutionary trajectory of this aquatic invertebrate.

## Materials and Methods

### Data acquisition

To examine the spatial distribution of mutations along the *C. riparius* genome, we reanalyzed published genomic datasets from five independent mutation‐accumulation related and exposure studies (Table 1).

**Table 1.** Datasets downloaded from ENA for the investigation of distribution and randomness of mutations along *C*.*riparius* genomes. Studies marked with * used an older version of the *C. riparius* genome. Ta = temperature, TRWPs= tyre and road wear particles and BaP= benzo[a]pyrene. Parenthesis indicate the original number of mutations reported in the study before the use of LiftOver to translate coordinates from old to new genome. The total cumulative number of mutations included in this study is 420.

| Factor | Number of mutations identified | Publication | ENA number |
| --- | --- | --- | --- |
| Spontaneous (T<sup>a</sup>)* | 14 (51) | Oppold and Pfenninger 2017 | PRJEB18039 |
| Cadmium exposure* | 66 (89) | Doria et al. 2021 | PRJEB40079 |
| TRWPs | 107 | Rigano et al. 2025 | PRJEB88551 |
| BaP exposure | 104 | Bulut et al. 2024 | PRJEB89420 |
| Generational time | 129 | Bulut and Pfenninger 2025 | PRJEB89193 |

**Table 2.** Bayesian Model Comparison using Leave-One-Out Information Criterion (LOOIC) and Expected Log Predictive Density (ELPD). The best-fitting model is in italics and listed first. ΔELPD is the difference in ELPD relative to the best model, with SE_diff_ denoting the standard error of this difference.

| Model | Hypothesis | LOOIC | ELPD | ΔELPD | $SE_{diff}$ |
| --- | --- | --- | --- | --- | --- |
| Model 4<br>($H_3$) | Dependence on distance to genes | 2244.6 | -1122.3 | 0.0 | 0.0 |
| Model 5<br>($H_4$) | Dependence on distance to repetitive elements | 2246.3 | -1123.1 | -0.8 | 0.2 |
| Model 3<br>($H_2$) | Dependence on distance to telomeres and centromeres | 2323.6 | -1161.8 | -39.5 | 6.7 |
| Model 2<br>(H<sub>1</sub>) | Non-random spatial<br>distribution | 2329.6 | -1164.8 | -42.5 | 7.3 |
| Model 1<br>(H<sub>0</sub>) | Random distribution | 2436.1 | -1218.1 | -95.8 | 6.4 |

For each dataset, we retrieved the list of single-nucleotide mutations (variant positions) in.bed format as reported in the respective studies. These genomic coordinates were used as input files for all downstream analyses. The chromosome scale *C. riparius* reference genome (Pettrich and Waldvogel 2025) served as the coordinate system for mapping and for the extraction of genomic feature information such as gene annotations, repeat elements, and chromosomal positions. Data from Oppold and Pfenninger (2017) and Doria et al. (2021) was obtained using the previous draft genome (Oppold et al. 2017). Therefore, it was necessary to translate the mutation positions reported in these studies to coordinates on the more recent genome assembly. In order to do so, we computed a pairwise genome alignment chains between the old (Oppold et al. 2017) and new *C*.*riparius* genome (Pettrich and Waldvogel 2025), using `lastz` (parameters K = 2400, L = 3000, Y = 9400, H = 2000, default scoring matrix), `axtChain` (default parameters except linearGap = loose), `RepeatFiller`, and `chainCleaner` (default parameters except minBrokenChainScore = 75,000 and -doPairs) (Suarez et al. 2017). We then used UCSC’s `liftOver` tool (Hinrichs et al. 2006) to lift the coordinates of mutation positions based on the old assembly to the new assembly.

### Bayesian modelling of mutation distribution

All analyses were conducted in R v4.3.2. To test alternative hypotheses about the spatial distribution of mutations, we applied a Bayesian modelling framework implemented in the `brms` package (Bürkner 2017). Data from all five studies were merged into a single dataset to increase statistical power and assess cumulative patterns across experimental conditions.

We tested five hypotheses: 1) H_0_ (null hypothesis) – random distribution: mutations occur randomly and independently along the genome; 2) H_1_ – non-random distribution: mutation density varies systematically across genomic regions; 3) H_2_ – mutation density varies with distance to telomeres and centromeres; 4) H_3_ – Mutation density varies with distance to genes and; 5) H_4_ – Mutation density varies with distance to repetitive elements. In order to test H_2_, H_3_ and H_4_, we used an in-house genome annotation to extract the position of centromeres/telomeres, genes and repetitive elements, respectively.

Each model was specified using the *brm()* function in `brms`, with mutation density per genomic window as the response variable and the relevant genomic covariate(s) as predictors. Appropriate priors were set for regression coefficients, and posterior sampling was carried out using four MCMC chains with 4,000 iterations each (1,000 warm-up). Model convergence was evaluated by inspecting trace plots and R-hat statistics (< 1.01 for all parameters).

Model comparison and selection were based on Leave-One-Out Cross-Validation (LOO-CV) using the *loo()* function, which estimates the expected log predictive density (ELPD) for each model. The model with the highest ELPD (and lowest ΔLOO) was considered best supported by the data. Posterior predictive checks were used to confirm adequate model fit and to visualize predicted versus observed mutation densities.

### Characterization of synonymous and non-synonymous mutations

After identifying the best-supported model, we further explored the functional composition of the detected mutations. Using `tbg-tools` v0.2 (https://github.com/philipp-schoennenbeck/tbg-tools; Schönnenbeck et al. 2021), we obtained counts of synonymous *vs* non-synonymous mutations occurring in exonic regions using the *C. riparius* reference genome annotation, and summary statistics were compiled across all datasets.

To evaluate whether 1) mutations are overrepresented in exons and, 2) the observed ratio of non-synonymous to synonymous mutations deviated from neutral expectations, we used the `BayesFactor` package in R (Morey and Rouder 2013) to test observed data *vs* neutral expectations. To address the first question, we first calculated the proportion of callable sites in the genome and proportion of callable sites in exonic regions, based on the genome assembly length and length of exonic regions based on the annotation. The proportion of callable sites in exonic regions was used as the neutral expectation. We tested two competing hypotheses: 1) H_0_ (null hypothesis): mutations occur at or below neutral expectation (neutral or depleted) and, H_1_: exonic mutations are enriched relative to random expectation. For the second question we tested two competing hypotheses: 1) H_0_ (null hypothesis): the observed proportion of non-synonymous mutations equals 0.75, consistent with predictions from neutral theory (Nei and Gojobori 1986) and, 2) H_1_: the observed proportion of non-synonymous mutations is higher than the predictions from neutral theory. The final count of non-synonymous mutations included in the second analysis excluded duplicated entries of transcripts for the same mutation position and 4-folded degenerate sites.

In both analyses the Bayes factor (BF_10_) was calculated to quantify the strength of evidence favoring H_1_ over H_0_. Following Jeffreys’ (1961) scale, values of BF_10_ > 3 were interpreted as moderate, > 10 as strong, and > 30 as very strong evidence for the alternative hypothesis. Posterior distributions and credible intervals were visualized to assess uncertainty around parameter estimates.

## Results

### Mutations distribution

A set of 420 mutations composed the final dataset for the downstream analysis. The original number of mutations from Oppold & Pfenninger (2017) and Doria et al., (2021), was reduced after the translation process from position from the draft to the most current *C. riparius* genome.

Results from the Bayesian framework analyses indicate that the null hypothesis of random mutation distribution (H_0_, Model 1) was strongly rejected. The intercept-only model demonstrated the poorest fit, with a LOOIC of 2436.1 (ELPD = -1218.1, SE = 46.4). All alternative hypotheses (H_1_–H_4_) provided substantially better fits to the data, confirming that mutation distribution is highly non-random (Figure 1a and Table 1).

**Figure 1.**
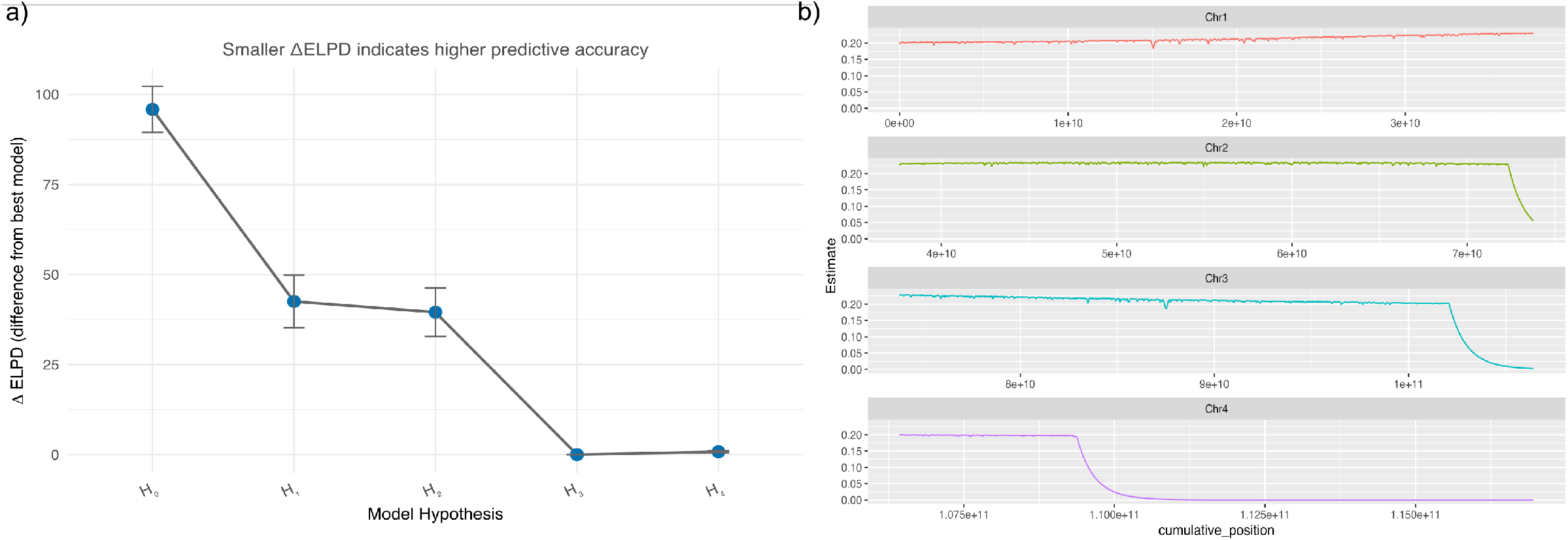
a) Model comparison using leave-one-out cross-validation (LOO). Points show the difference in expected log predictive density (ΔELPD) relative to the best model; horizontal bars are ± one standard error of the difference. Positive values indicate better predictive performance. Pareto *k* diagnostics and full LOOIC /ELPD values are provided in Table 1; b) Predicted mutation rate across the genome under the best-supported Bayesian model (Model 4). Posterior mean estimates (solid lines) and 95% credible intervals (shaded areas) are shown for each chromosome, based on the inhomogeneous Poisson process fitted in *brms* using genomic position and distance to the nearest gene as smooth predictors. The model captures spatial variation in mutation rate along each chromosome, with a largely uniform rate across most regions and a marked decline toward chromosomal ends, particularly on chromosomes 3 and 4.

**Figure 2.**
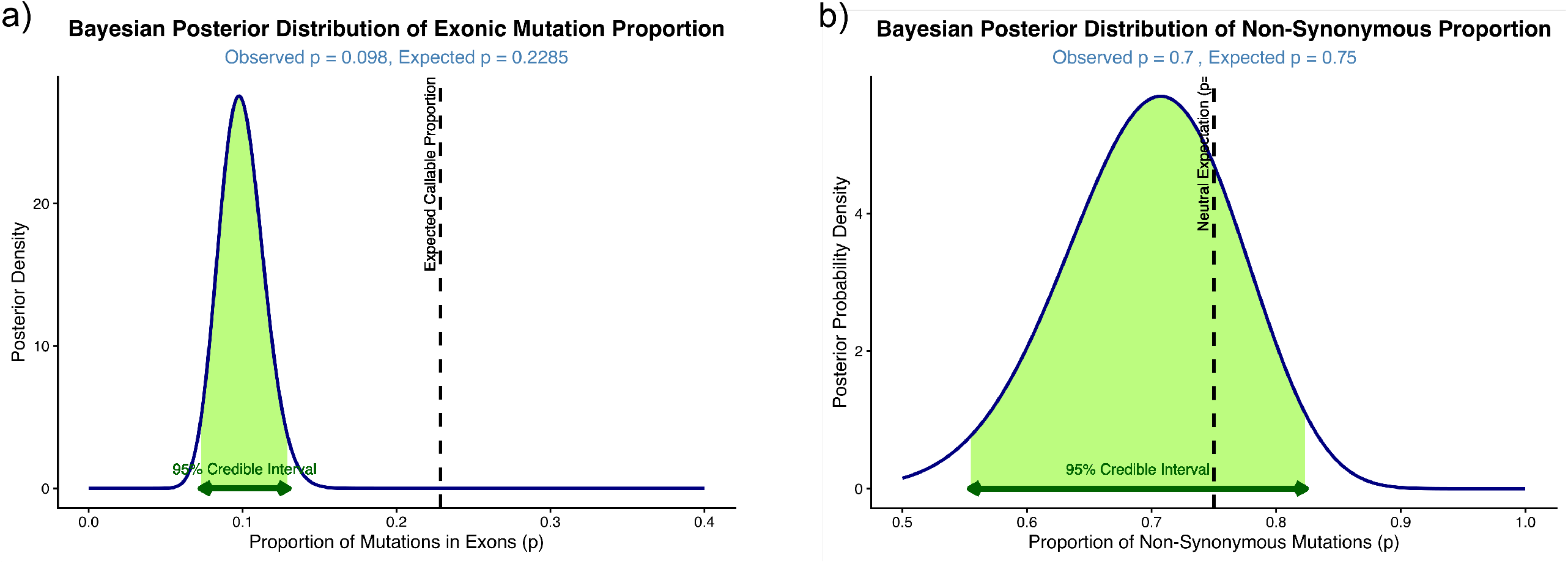
Posterior probabilities distributions. a) indicates the expected proportion of occurring in exons (green curve) *vs* the observed proportion of mutations in exons (dash line) and; b) indicates the expected proportion of non-synonymous mutation (green curve) *vs* the observed proportion of non-synonymous mutations (dash line).

The model comparison clearly identified Model 4 (H_3_: dependence on distance to genes) as the best-fitting model, exhibiting the highest ELPD (or lowest LOOIC) value (LOOIC = 2244.6, ELPD = -1122.3, SE = 42.0). The next-best model, Model 5 (H_4_: distance to repetitive elements), showed a slightly worse fit, with a difference in ELPD of ΔELPD = -0.8 (SE_diff_ = 0.2) compared to the top model. Models incorporating only genomic position or distance to telomeres/centromeres (Models 2 and 3) were significantly worse than the best model (e.g., ΔELPD_Model 3_ = -39.5, SE_diff_ = 6.7).

The best-fitting model (Model 4), which included a non-linear term for cumulative position and distance to genes, confirmed a strong relationship between mutation count and proximity to genes. The smoothing term for distance to genes was highly informative, with its posterior mean estimate indicating a substantial non-linear effect that was clearly distinguishable from zero (Estimate = -43.10, 95% CI = [-66.54, -23.06]). This suggests that mutation accumulation is significantly modulated by proximity to gene features. In contrast, the smoothing term for the genome-wide cumulative position, which accounts for broad spatial effects, did not show strong evidence for a positional effect once distance to genes was accounted for (Estimate = -0.07, 95%CI = [-3.16, 3.08]).

### Functional enrichment of mutation in *C. riparius*

Of 420 total mutations, 41 (9.8%) occurred in exonic regions. Given that 22.85% of callable sites in the genome are exonic, this proportion is markedly lower than expected under a random distribution. A Bayesian proportion test (H_1_: p > 0.2285 *vs* H_0_: p ≤ 0.2285) yielded a Bayes Factor of ≈ 97, providing very strong evidence for the null hypothesis and indicating a significant depletion of exonic mutations. The true proportion of mutations occurring in exons lies outside 95% CI (95% CI: [0.073, 0.130] (Figure 2a).

Of 41 exonic mutations identified by tbg-tools, 29 (70.7%) were non-synonymous. A Bayesian proportion test (H_1_: p > 0.75 *vs* H_0_: p ≤ 0.75) yielded BF_10_ ≈ 31, providing very strong evidence in favor of the null hypothesis. The observed proportion of non-synonymous mutations does not exceed neutral expectations. The true proportion of non-synonymous mutations lies between the 95% CI (95% CI : [0.554, 0.824]) (Figure 2b).

## Discussion

Our results provide evidence that mutations in the genome of *C. riparius* are not distributed randomly. By explicitly testing multiple competing hypotheses within a Bayesian framework, we demonstrate that the null model assuming spatial randomness (H_0_) is strongly rejected. Instead, mutation accumulation is significantly modulated by proximity to genes, supporting the hypothesis that genomic architecture exerts a strong influence on local mutation rates. Moreover, although the overall mutation rate is heterogeneous, exonic regions are markedly depleted of mutations, and the proportion of non-synonymous changes among those mutations conforms to neutral expectations. Taken together, these findings indicate that mutation rate variation in *C. riparius* is primarily structured by the genome’s functional organization rather than being a product of stochastic mutational noise. This study contributes to the growing body of evidence that the distribution of mutations within eukaryotic genomes is influenced by both structural and functional constraints (Hodgkinson and Eyre-Walker 2011). By focusing on *C. riparius*—a non-model aquatic insect of ecological importance—we extend this framework to a taxon that experiences strong environmental stress and has a relatively compact but repeat-rich genome. Our results suggest that even in such systems, mutation processes are neither uniform nor purely random, but rather reflect the interplay between genome architecture, DNA repair efficiency, and functional constraint.

### Evidence for non-random mutation patterns

The rejection of the random-distribution model (H_0_) provides the first genome-wide statistical evidence that *C. riparius* mutation accumulation deviates significantly from a homogeneous Poisson process. In classical population genetics, randomness in mutation implies that the expected number of mutations per genomic window follows a constant rate across the genome, with variance proportional to the mean. However, our data exhibit significant overdispersion, and model comparison using the Leave-One-Out Information Criterion (LOOIC) clearly identified spatial and functional predictors as superior explanatory variables.

Models incorporating genomic features (H_1_–H_4_) substantially improved predictive power, but the most informative model (H_3_) revealed that distance to genes was the dominant predictor of mutation density. This implies that mutation rate heterogeneity in *C. riparius* is tightly coupled to the genome’s functional landscape. The second-best model (H_4_), based on proximity to repetitive elements, provided a nearly equivalent fit (ΔELPD = –0.8), suggesting that both functional (genic) and structural (repetitive) features jointly shape the mutational landscape. In contrast, models based on chromosome-scale effects such as distance to telomeres or centromeres (H_2_) contributed little explanatory power once gene proximity was considered.

This pattern aligns with previous studies in mammals, plants, and *Drosophila*, which have shown that genes and nearby regions exhibit reduced mutation rates relative to intergenic DNA (Bzymek and Lovett 2001; Polak et al. 2014). The mechanistic basis for this pattern is often linked to transcription-coupled repair (TCR), which preferentially removes lesions from the template strand of actively transcribed genes (Deger et al. 2019; Törmä et al. 2020). Our results suggest that a similar repair bias is present in *C. riparius*, resulting in lower mutational input near genic regions. Because *C. riparius* is frequently exposed to environmental mutagens, the efficiency of local repair processes may play a crucial role in maintaining genomic integrity under stress.

The best-supported model revealed a strong negative effect of distance to genes on mutation density (posterior estimate = –43.1; 95% CI = [–66.5, –23.1]). This nonlinear pattern suggests that the mutational landscape transitions sharply from low to high mutation density with increasing distance from coding regions. Such a gradient is consistent with the hypothesis that repair processes, chromatin openness, and replication timing create a local shield around genes.

In eukaryotic genomes, the chromatin state strongly influences the accessibility of DNA repair enzymes. Gene-rich regions are often euchromatic, transcriptionally active, and associated with high-fidelity repair mechanisms such as homologous recombination and TCR. In contrast, intergenic and heterochromatic regions—where *C. riparius* accumulates most of its mutations— are subject to less efficient, error-prone repair pathways such as NHEJ (Chen and Tyler 2022). This spatial segregation of repair processes can explain the strong decay of mutation rate with distance from genes observed in our data.

In several species, however, regions immediately adjacent to highly expressed genes can exhibit locally elevated mutation rates due to transcription-associated mutagenesis (Polak et al. 2014). The smooth, nonlinear term in our model likely captures this complexity—indicating that while genes themselves are protected, the flanking regions might occasionally act as “mutation buffers” that absorb transcription-induced damage without compromising essential coding sequences.

Although the gene-distance model (H_3_) performed best, the model incorporating proximity to repetitive elements (H_4_) displayed nearly equivalent explanatory power. This suggests that repetitive DNA also contributes significantly to mutation rate variation in *C. riparius*. The genome of this species contains a high density of repetitive and transposable elements (Oppold et al. 2017; Pettrich and Waldvogel 2025), which are inherently unstable due to replication slippage and recombination events (Bzymek and Lovett 2001; Ellegren 2004; Cabral-de-Mello and Palacios-Gimenez 2025).

Repetitive elements can promote local genome instability through several mechanisms: (1) mispairing during replication, (2) unequal crossing-over during recombination, and (3) formation of secondary structures such as hairpins that interfere with polymerase progression. In addition, transposable element (TE) insertions can locally alter chromatin structure, potentially disrupting nearby DNA-repair dynamics and generating localized mutational hotspots (Bzymek and Lovett 2001). The near equivalence of models H_3_ and H_4_ therefore reflects an interplay between functional (genic) and structural (repetitive) influences on mutation probability.

The strong spatial heterogeneity in mutation accumulation observed here is consistent with the concept of MRH described in various taxa (Seplyarskiy et al. 2012; Liu et al. 2017; Nesta et al. 2021). In *C. riparius*, MRH appears to arise from both intrinsic genomic architecture and extrinsic environmental factors—since this aquatic species is constantly exposed to mutagens such as heavy metals or organic pollutants (Doria et al. 2021; Bulut et al. 2024; Rigano et al. 2025). These results thus bridge molecular evolutionary theory with environmental genomics, demonstrating that environmental mutagenicity acts on an already heterogeneous genomic template.

### Non-synonymous mutations and the signature of selection

Our analyses revealed a striking depletion of mutations in exonic regions. Only 9.8% of all detected mutations were exonic, even though 22.85% of callable genomic sites correspond to exonic sequences. The Bayesian test yielded a Bayes Factor (BF_10_ ≈ 97), providing very strong support for the null hypothesis that exonic mutations occur less frequently than expected under random distribution. The 95% credible interval ([0.073, 0.130]) further confirms this depletion.

This finding aligns with results from other eukaryotic systems where exons display lower mutation rates than intergenic or intronic DNA (Subramanian and Kumar 2003; Frigola et al. 2017; Rodriguez-Galindo et al. 2020). Multiple non-mutually exclusive mechanisms can account for this pattern: 1) Enhanced DNA repair efficiency in coding regions. Genes, particularly those that are actively transcribed, benefit from transcription-coupled repair and higher surveillance by DNA repair enzymes. This reduces the fixation probability of spontaneous or environmentally induced mutations within exons; 2) Purifying selection acting on deleterious alleles. Even in mutation-accumulation or short-term experimental contexts, highly deleterious exonic mutations can be rapidly purged from germline lineages, especially in large effective populations. This may reduce the observed number of exonic variants relative to intergenic sites and; 3) Chromatin context and replication timing. Early-replicating euchromatic regions, which are enriched in genes, tend to exhibit lower mutation rates due to more accurate replication and efficient error correction (Moore et al. 2021).

In *C. riparius*, the strong depletion of exonic mutations is particularly noteworthy given the organism’s exposure to genotoxic pollutants. It suggests that despite increased environmental mutagenesis, intrinsic genomic and cellular mechanisms remain efficient in safeguarding coding integrity—a finding consistent with the adaptive resilience expected from a species occupying highly variable aquatic environments.

Additionally, of the 41 exonic mutations, 29 (70.7%) were non-synonymous. The Bayesian proportion test yielded a Bayes Factor of ≈31, providing strong evidence against enrichment beyond the neutral expectation (p = 0.75). The posterior credible interval ([0.554, 0.824]) overlapped substantially with the neutral benchmark. This indicates that, among exonic mutations that do occur, the ratio of amino-acid–changing to silent mutations conforms to theoretical expectations derived from the genetic code structure (Nei and Gojobori 1986). This result has several important implications. First, it suggests that the mutational process itself— once constrained to exonic sites—is not biased toward producing a higher proportion of functionally impactful changes. The depletion of exonic mutations observed earlier thus likely reflects where mutations occur, rather than how the genetic code is altered once a mutation happens. In other words, *C. riparius* does not exhibit an unusual codon-level mutation bias; rather, its genome architecture restricts the occurrence of mutations in coding regions altogether. Second, the consistency with neutral expectations implies that the majority of mutations captured in our dataset arose under conditions with limited selective filtering—consistent with the mutation-accumulation and exposure-based designs of the original datasets (Oppold and Pfenninger 2017; Bulut et al. 2024). Thus, even though environmental stress can elevate mutation rates, it does not necessarily skew the spectrum of synonymous *vs* non-synonymous changes, at least within the resolution of the available data.

### Evolutionary and ecotoxicological implications for *C. riparius*

The fine-scale understanding of the mutation landscape in *C. riparius* has significant implications for both evolutionary genomics and ecotoxicology. Our data reveal a strongly asymmetric mutational landscape: genes are effectively shielded from mutation, but repetitive regions—where repair efficiency appears lower—exhibit substantially elevated mutation rates. These repeat-rich regions therefore constitute the major contributors to the mutational load in *C. riparius*.

The pronounced influence of gene proximity and the depletion of exonic mutations point to heterogeneous DNA repair efficiency as a key driver of mutation rate variation. Similar genomic gradients have been reported in humans and *Drosophila*, where transcription-coupled and base-excision repair pathways preferentially protect genic regions (Supek and Lehner 2015; Glastad et al. 2015). In *C. riparius*, these processes may be further influenced by environmental stressors that either induce or compromise specific repair mechanisms.

The ecology of *C. riparius* adds an environmental dimension to this pattern. Living in sediment-rich aquatic habitats, the species is chronically exposed to heavy metals, hydrocarbons, and microplastic-derived pollutants that induce oxidative and alkylating DNA damage (Doria et al. 2021; Rigano et al. 2025). Because such damage is repaired unevenly across the genome, environmental stress likely amplifies pre-existing mutation rate heterogeneity (MRH). Our findings suggest that *C. riparius* has evolved structural defenses—such as enhanced repair near essential loci—that mitigate damage in functional regions while tolerating higher variability in repeat-rich areas. This pattern supports conceptual models proposing that MRH itself can be adaptive, buffering vital genes from deleterious change while maintaining genome-wide evolvability (Seplyarskiy et al. 2012; Liu et al. 2017). In this light, the mutation landscape of *C. riparius* may represent an evolved compromise between genomic stability and flexibility under persistent mutagenic stress.

From a population genetics perspective, the strong spatial heterogeneity in mutation rate means that standard models assuming uniform mutagenesis are inadequate for *C. riparius*. Regions with elevated mutation density—typically gene-poor intergenic DNA—may exhibit inflated polymorphism levels, biasing estimates of effective population size (N_e_), divergence times, or introgression if not properly accounted for. Future analyses should therefore include genomic position as a covariate when estimating mutation-dependent parameters. Mutation rate heterogeneity also contributes directly to differences in nucleotide diversity and substitution rates across the genome (Hodgkinson and Eyre-Walker 2011). Gene-rich, repair-efficient regions are expected to conserve ancestral states, whereas repetitive regions accumulate neutral or slightly deleterious variants that may later contribute to adaptation. In *C. riparius*, which exhibits high connectivity and gene flow (Waldvogel et al. 2018), heterogeneous mutation input could interact with recombination and selection to shape local adaptation. If mutation hotspots overlap with regions under pollutant-driven selection, they might serve as reservoirs for adaptive variation.

As a standard ecotoxicological model, *C. riparius* is pivotal for understanding the genomic consequences of environmental contamination. Exposure to complex pollutant mixtures, such as tyre and road wear particles (Rigano et al. 2025), has been shown to elevate germline mutation rates. Knowing where these new mutations occur is crucial. Our results suggest that while genes are protected by efficient repair, most induced mutations accumulate in gene-poor, highly mutable regions. This “dark matter” polymorphism in intergenic DNA could still affect regulatory sequences, chromatin organization, or transposable element dynamics, influencing long-term genome evolution even if coding regions remain buffered.

Future work should test whether chronic pollutant exposure alters the spatial distribution of mutations by suppressing transcription-coupled repair (TCR) and global genome repair (GGR), or merely increases the baseline rate of mutation across the genome. Controlled experiments combining mutation accumulation, transcriptomics, and repair-pathway profiling could clarify these mechanisms. Integrating such insights with environmental monitoring would establish stronger mechanistic links between pollutant exposure, DNA repair capacity, and genomic mutability, bridging molecular evolution with ecological toxicology.

## Conclusions

This study contributes to the growing body of evidence that the distribution of mutations within eukaryotic genomes is influenced by both structural and functional constraints. By focusing on *C. riparius*—a non-model aquatic insect of ecological importance—we extend this framework to a taxon that experiences strong environmental stress and has a relatively compact but repeat-rich genome. Our results suggest that even in such systems, mutation processes are neither uniform nor purely random, but rather reflect the interplay between genome architecture, DNA repair efficiency, and functional constraint. Our study provides clear evidence that the distribution of mutations in the *C. riparius* genome is non-random and shaped by the underlying genomic architecture. Mutation accumulation is significantly modulated by proximity to genes and, to a lesser extent, by the presence of repetitive elements. Coding regions are strongly protected from mutational input, consistent with efficient repair and purifying selection, while the relative frequency of non-synonymous mutations conforms to neutral expectations once such constraints are accounted for.

These results underscore the importance of considering structural and functional genomic context in models of mutation rate and evolutionary potential. In *C. riparius*, the mutational landscape appears to balance genome stability with flexibility—protecting functionally essential loci while permitting variability in less constrained regions. Such a balance may be particularly adaptive for species inhabiting environments with high and variable mutagenic stress.

By combining comparative genomics with Bayesian spatial modelling, our study bridges molecular, ecological, and evolutionary perspectives on mutation processes. It demonstrates that even in a small insect genome, mutation is far from random—reflecting the intricate interaction between genome architecture, repair mechanisms, and environmental context that ultimately shapes the pace and direction of evolution.

## Acknowledgements

We thank Dr. Tilman Schell and Prof. Dr. Michael Hiller for their advice and assistance in the use of `liftOver`.

## Data Availability Statement

The analyses presented here are based on the mutation lists provided as supplementary Information in the five publications used (Table 1). The original raw sequencing data archived in ENA was not used.

